# RNA-controlled nucleocytoplasmic shuttling of mRNA decay factors regulates mRNA synthesis and initiates a novel mRNA decay pathway

**DOI:** 10.1101/2021.04.01.437949

**Authors:** Shiladitya Chattopadhyay, Jose Garcia-Martinez, Gal Haimovich, Aya Khwaja, Oren Barkai, Ambarnil Ghosh, Silvia Gabriela Chuarzman, Maya Schuldiner, Ron Elran, Miriam Rosenberg, Katherine Bohnsack, Markus Bohnsack, Jose E Perez-Ortin, Mordechai Choder

## Abstract

mRNA level is controlled by factors that mediate both mRNA synthesis and decay, including the exonuclease Xrn1 - a major mRNA synthesis and decay factor. Here we show that nucleocytoplasmic shuttling of Xrn1 and of some of its associated mRNA decay factors plays a key role in determining both mRNA synthesis and decay. Shuttling is regulated by RNA-controlled binding of the karyopherin Kap120 to two nuclear localization sequences (NLSs) in Xrn1. The decaying RNA binds and masks NLS1, establishing a link between mRNA decay and Xrn1 shuttling. Mutations in the two NLSs, which prevent Xrn1 import, compromise transcription and, unexpectedly, also the cytoplasmic decay of ∼50% of the cellular mRNAs - comparably to Xrn1 deletion. These findings uncover a cytoplasmic mRNA decay pathway that begins in the nucleus. Interestingly, Xrn1 shuttling is required for proper adaptation to environmental changes, in particular to ever changing environmental fluctuations.

## Main

A high degree of regulation of mRNA levels is a critical feature of gene expression in any living organism. In recent years we and other investigators have discovered reciprocal adjustments between the overall rates of mRNA synthesis and degradation, named “mRNA buffering”, which maintain proper concentrations of mRNAs^1–7^. We have previously demonstrated that in the budding yeast *Saccharomyces cerevisiae* (from herein termed yeast), a number of factors, known to degrade mRNA in the cytoplasm, e.g., Xrn1 (alias – Kem1), Dcp2, Pat1, Lsm1^8, 9^, shuttle between the nucleus and the cytoplasm, by an unknown mechanism^3^. The mRNA buffering mechanism is not restricted to factors recognized as mRNA decay factors (DFs). It also includes components of the transcription apparatus. We demonstrated that Pol II regulates mRNA translation and decay by mediating Rpb4/7 co-transcriptional binding to Pol II transcripts^10–12^, a process we named “mRNA imprinting”^13, 14^. More “classical” yeast DFs – components of the Ccr4-Not complex - also imprint mRNA and regulate mRNA export, translation and decay ^15, 16^. Thus, cellular localization of these factors seems to be critical for their two opposing activities, either in synthesis or degradation.

Here we show that Xrn1 nuclear import, mediated by two NLSs and by the decaying RNA, affects mRNA synthesis and decay of a large portion of the mRNAs. We define the affected mRNAs as Kem1 import-sensitive (Kis) mRNAs. Half-lives (HLs) of Kis mRNAs are affected by blocking Xrn1 import, indicating that the nuclear function of Xrn1 is indispensable for Xrn1-mediated cytoplasmic decay. We propose that the decay pathway of Kis mRNAs begins in the nucleus, probably by Xrn1 binding to these mRNAs. Thus, our study unveils a nuclear step of the cytoplasmic decay of ∼50% of the mRNAs. Our model proposes that many mRNAs are exported to the cytoplasm while already carrying some of the degradation factors with them. It suggests that transcription has a much stronger effect on mRNA decay than we have viewed thus far.

## Results

### Xrn1 contains two functional nuclear localization sequences (NLSs)

Using cNLS mapper^17–19^ and NLStradamus^20^ programs we found two potential NLSs in Xrn1, but not in other DFs. NLS1 is a classical monopartite NLS (368-LEGERKRQRVGK-381), located at the junction between the “core” enzyme and a “linker domain”^21^. This linker, whose function is unknown, protrudes outside of the core and will be referred herein as “Tail” (Fig. 1A). NLS1 sequence and its location within Xrn1 structure are conserved in different yeast species (Fig. 1B). Presence of NLS in a tail-like protrusion of Xrn1 was also found in Xrn1 homologues in higher eukaryotes (Fig. S1A-B). NLS2 (1246-KKALEKKK-1255), which is located in the C-terminus (Fig. 1A), does not show significant sequence conservation; nevertheless, basic NLSs are present around the same amino-acid position in different yeast strains (Fig. S1C). Both NLSs are located in intrinsically disordered regions (Fig. S1D). Using an import assay^3, 22^, we found that the two NLSs are functional, as replacing the 6 basic residues of NLS1 with alanine (ΔNLS1) compromised Xrn1-GFP import (Fig. 1C-D and S1E); Likewise, replacing 4 basic residues of NLS2 with alanine (ΔNLS2) decreased import (Fig. 1C-D and S1E). Import of Xrn1-GFP carrying the mutations in both NLSs (Xrn1^ΔNLS1/2^-GFP) is almost entirely blocked (Fig. 1C, S1E) while that of a co-expressed Pab1-RFP was unaffected by the mutations (Fig. 1D). A single deletion of any of them affected Xrn1 import (Fig. 1C, Fig. S1E).

**Fig. 1.**
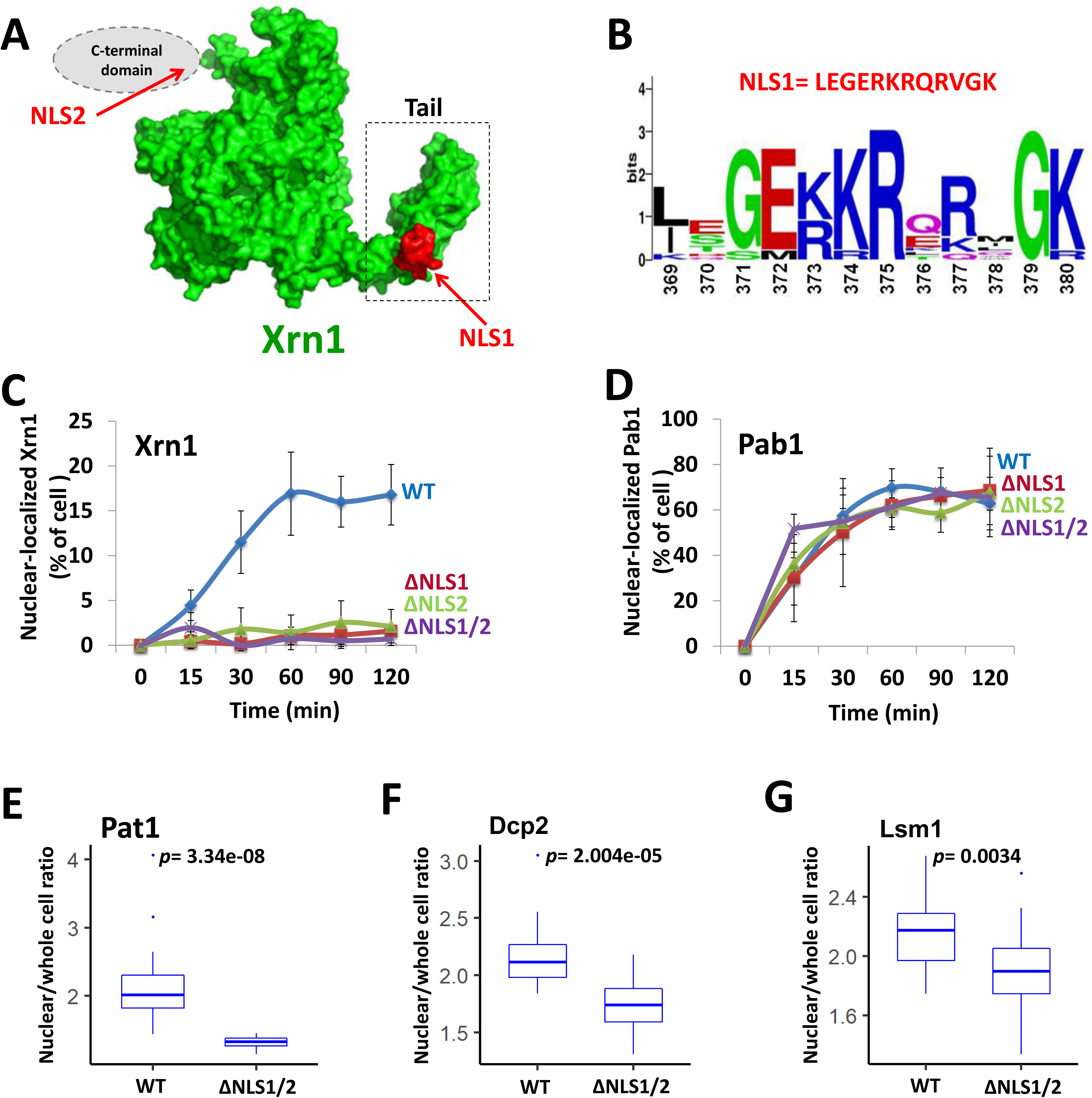
Xrn1 contains two functional NLSs. (**A**) *Position of NLSs*. NLS1, found by cNLSmapper, is highlighted in red. Xrn1 structure was modelled using SWISS-MODEL^63^. Tail is indicated by a dashed box. The expected location of the unstructured C-terminal domain containing NLS2 is shown schematically as a grey ellipse. (**B**) *Sequence conservation of NLS1 among different yeasts*. The sequences of 11 yeast stains were compared by the online WebLogo (U. Berkeley). The height of each stack of letters represents the degree of sequence conservation, measured in bits. (**C**) *Xrn1-GFP import kinetics*. A standard import assay was performed, as described in Methods, using cells that co-expressed Pab1p-RFP (served as positive control, see D) and Xrn1-GFP or its mutant derivatives, as indicated. Percentage of cells showing nuclear import was measured at the indicated time intervals after inactivation of protein export. (**D**) *Import of Pab1, used as a positive control*. Error bars in C and D represent standard deviation (S.D.) of three independent assays. (**E-G**) *Starvation-induced nuclear localization of the indicated fluorescent DFs is dependent on Xrn1 NLSs*. WT or *xrn1^Δ^*^NLS1/2^ cells expressing the indicated GFP fused DF were shifted from optimal conditions to a starvation medium lacking sugar and amino acids. After 1 h, the fluorescence distribution was measured using ImageJ. The image segmentation, based on Rpb4-GFP as the nuclear marker, was done by thresholding and selecting area of the nucleus. The Mean Fluorescence intensity of the nucleus was thus determined. Three replicates were used, 5-8 fields (>300 cells) per replicate. The ratio between the nuclear and whole-cell fluorescence is shown as a boxplot. *p*-value was calculated by Wilcoxon rank sum test.

We have previously reported that several DFs are imported to the nucleus and function together in transcription, probably as a complex^2,3^. To determine whether import of Pat1, Dcp2 and Lsm1 is dependent on Xrn1, we took advantage of a previous observation that during nutrient starvation they are found in the nucleus^3^. Indeed, in xrn1^ΔNLS1/2^ cells these DFs remain cytoplasmic during starvation (Fig. 1E-G and S1F-H). Thus, Xrn1 NLSs are involved, directly or indirectly, in importing also other DFs.

### Identification of mutations that cause constitutive nuclear localization of Tail revealed an unidentified RNA binding domain

To further study NLS1 function outside the context of Xrn1 and NLS2, we replaced the Xrn1 core sequences with GFP, creatinga “Tail-GFP” fusion protein. Unexpectedly, despite the presence of NLS1, GFP-tagged Tail localized in the cytoplasm (Fig. 2A). However, the Tail-GFP can be imported (Fig. 2B), indicating that Tail-GFP shuttles between the two compartments. We conclude that its apparent exclusion from the nucleus is due to its slow import relative to its export kinetics.

**Fig. 2.**
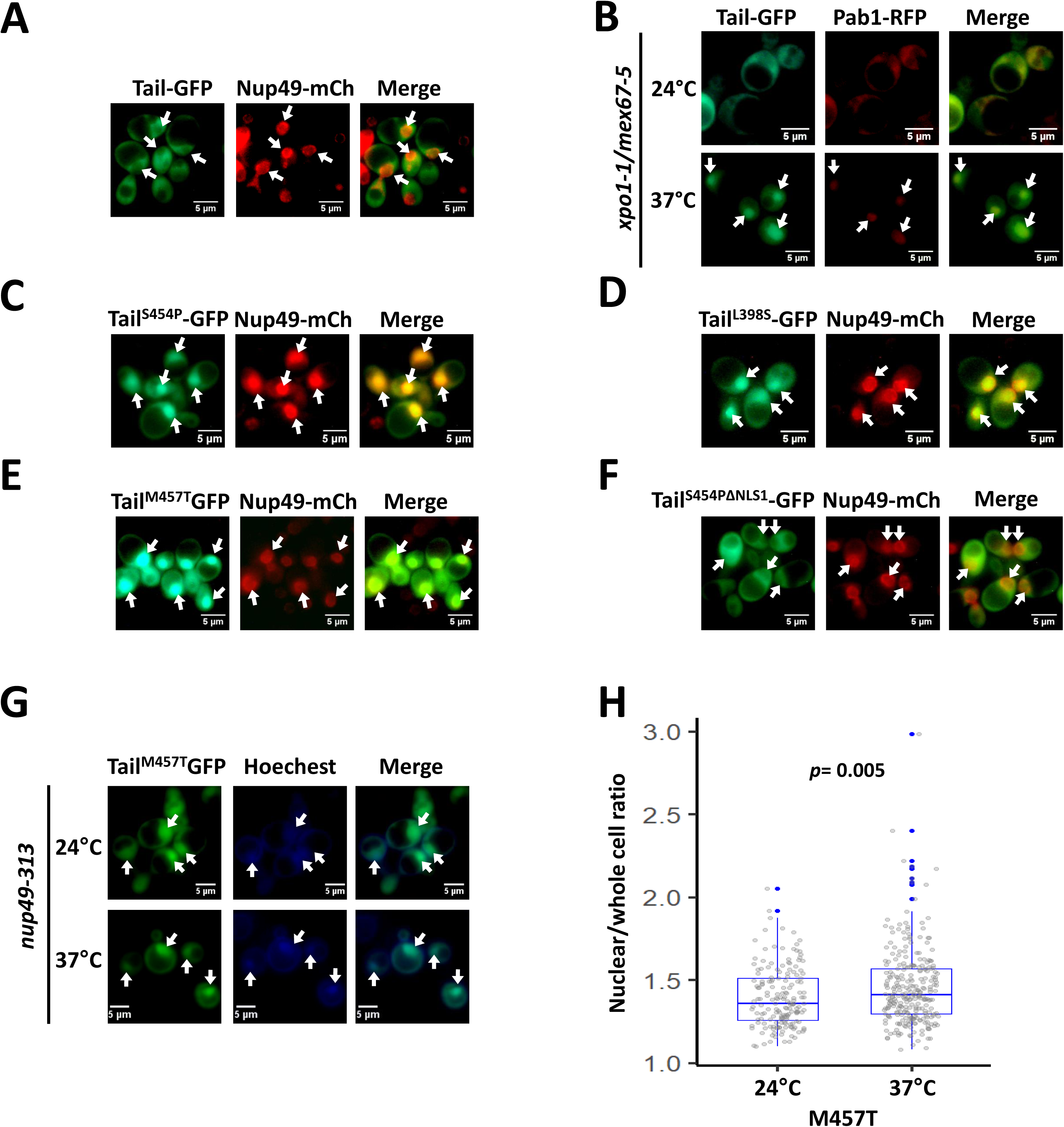
Some mutant Tail-GFP molecules are localized constitutively in the nucleus. Tail (amino acid 351 until 521 of the *XRN1* sequence) was fused to GFP and was expressed under *ADH1* promoter and terminator. Nup49-RFP (a nucleoporin) was used as a nuclear marker. (**A**) *During optimal proliferation Tail is mainly cytoplasmic*. (**B**) *Tail-GFP shuttles between cytoplasm and nucleus*. Tail-GFP localization was examined by the import assay, as in Fig. 1C. Photos were taken after 2h at 37°C. (**C-E**) *Mutant Tail-GFP molecules that are constitutively localized to the nucleus*. The Tail sequence of Tail-GFP was randomly mutagenized by PCR mutagenesis protocol and introduced into WT strain by transformation. Distinct colonies were allowed to proliferate in 96 wells. The cellular localization of the mutants was automatically scanned by fluorescent microscope using a robot. Shown are a few of the mutants that were localized to the nucleus in optimally proliferating cells. See Fig. S2D for mapping of mutations. (**F**) *Tail^S454P^ nuclear import is mediated by NLS1*. NLS1 of Tail^S454P^ was mutagenized (resulting in Tail^S454P, ΔNLS1^), and the mutant was analyzed as in C. (**G**) *Localization of mutant Tail-GFP in temperature-sensitive (ts) nuclear-import mutant*. A shuttling assay of Tail^M457T^ in the ts *nup49-313* import mutant was performed as described previously^67^. (**H**) *Quantification of the shuttling assay*. The nuclear/whole-cell ratio of the mean fluorescence intensity was determined by ImageJ, as described in Methods. P-value, based on 3 replicates, was calculated by Wilcoxon rank sum test (H).

After randomly mutagenizing Tail, a number of mutants were identified to localize almost exclusively in the nucleus (except for R507G) (Fig.2C-E, quantification of the nuclear/whole-cell ratio of the mean fluorescence intensity is shown in Fig. 4E, WT). Thus, these mutations either compromised Tail export or enhance its import kinetics or both. Most of the single mutations map to a structural motif, that resembles a known RNA-binding motif R3H (Fig. S2). The RNA binding potential of the tail is also supported by recently published cryo-EM structure of Xrn1 bound to ribosomal RNA ^23^. Tail^S454P^-GFP is one of the constitutive nuclear mutants (Fig. 2C), but disruption of its NLS1 blocked its import (Fig. 2F), indicating that NLS1 is necessary for importing Tail-GFP. Both Tail^S454P^-GFP and Tail^M457T^-GFP remain in the nucleus upon heat inactivation of the nucleoporin Nup49-313 and repression of protein import (Fig. 2G-H) ^22, 24–26^, suggesting that these mutations compromised Tail export.

### Tail binds RNA via an R3H-like motif

We determined the RNA-binding feature of HTP-tagged Tail by UV cross-linking cells followed by Tail affinity purification, RNA fragmentation and labelling. SDS-PAGE of radioactive immunoprecipitated (IPed) material confirmed that Tail can bind RNA when expressed outside Xrn1 context (Fig. 3A). RNA binding of Tail^S454P^ and Tail^M457T^, carrying mutations that also caused constitutive nuclear localization (Fig. 2C and E), was weaker than the WT tail (Fig. 3B), indicating that at least two mutations that affect the shuttling feature of the tail and are located in the R3H-like motif also affect RNA-binding.

**Fig.3.**
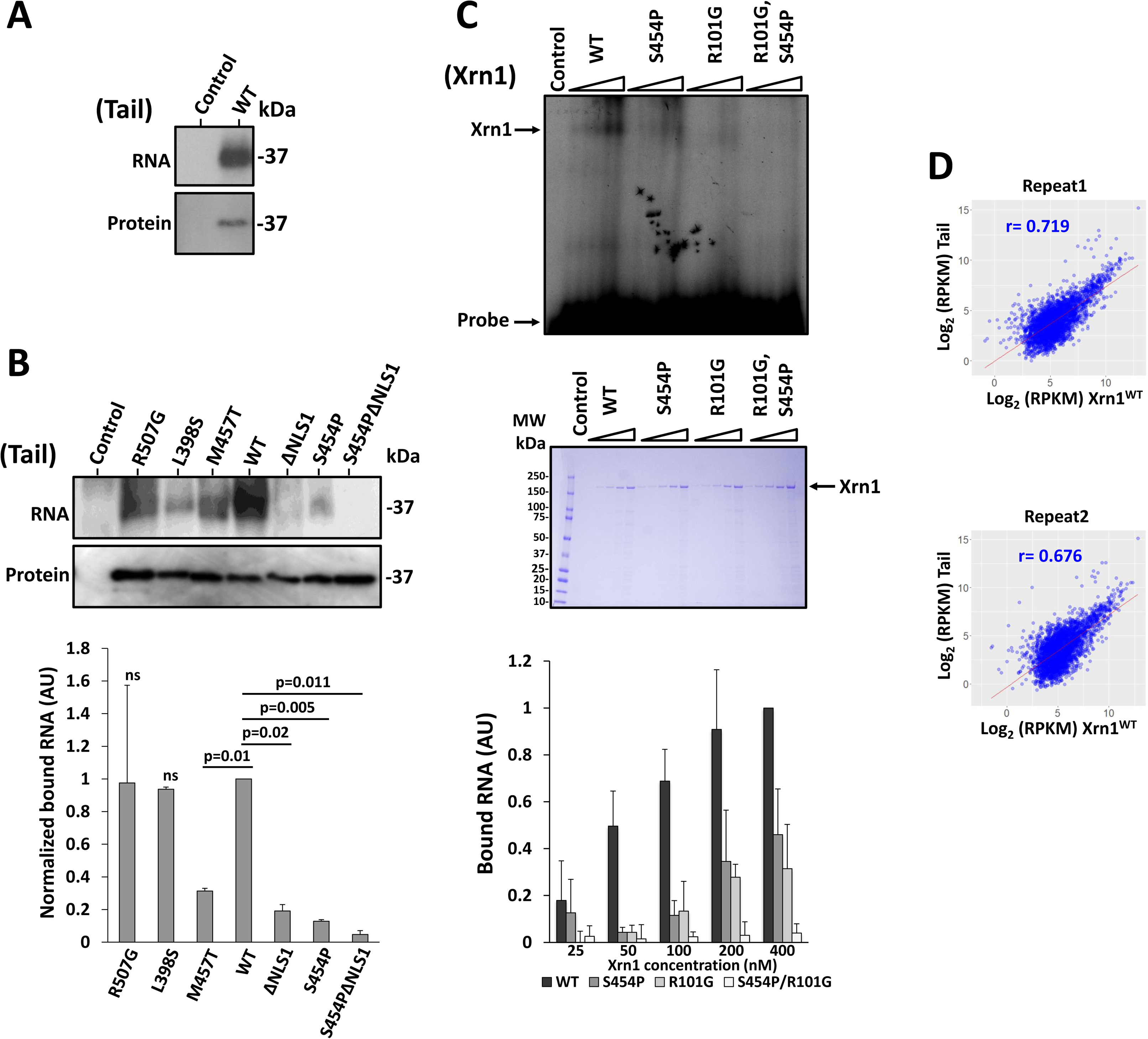
Xrn1 Tail binds RNA. (**A**) *Tail binds RNA*. Cells expressing 6xHis-TEV cleavage site-Protein A (HTP) tagged Tail, or control cells carrying no tag,were UV crosslinked. Tail-HTP-RNA complex was affinity purified, labelled and run on SDS gel as described in Methods. Upper panel, autoradiogram (RNA); bottom panel, Western blot using anti-HISx6 antibodies to detect the protein. (**B**) *Some mutations in Tail (indicated above each lane) affect its RNA binding capacity*. Affinity purification of the indicated Tail mutants, which had been crosslinked to RNA, was performed as in A. The RNA and protein levels were quantified as described in Methods. RNA levels were normalized to the protein level. Results represent mean (±S.D.) of 2 replicates. WT is arbitrarily defined as 1. Statistical significance was determined by unpaired t-test and *p*-values are indicated. (**C**) *Tail provides a major RNA binding site of Xrn1 which acts in conjunction with the R101 residue of the active site*. FLAG-tagged Xrn1, or it’s indicated mutant derivatives, were affinity purified (see Methods), and subject to a gel-shift assay using ^32^P-5’-RNA (a mixture of 40, 32 and 28 b long), in the presence of EDTA to inactivate the enzyme (see Methods). Increasing amounts of purified proteins were used as indicated. Control lane represents experiment in which the probe was reacted with elute from untagged cells. Quantification of 3 replicates (mean ± S.D.) is shown at the bottom. The results of WT Xrn1 reacted with 400 µM was defined arbitrarily as 1. Purified Xrn1 was run in parallel, to verify equal amount of protein loading. Shown is a Coomassie blue straining (middle panel). (**D**) *RNA bound by Tail is highly correlated to Xrn1 RNA interactome*. Tail and Xrn1 were subject to a full CRAC analysis. RPKM values of RNA binding to Tail and Xrn1 were scatterplotted. Two replicates are shown. Pearson’s correlation (r) is indicated.

Since NLS1 is positively charged and closely located to the RNA-binding motif, we surmised that NLS1 binds RNA as well, and hypothesized that this binding masks NLS1 from recognition by potential importins. Indeed, RNA-binding capacity of the Tail^ΔNLS1^ was defective (Fig. 3B), indicating that NLS1 also contributes to the RNA binding feature of the Tail. RNA binding of Tail carrying mutations in both S454P and NLS1 was 22-fold weaker than that of WT and 2.7-fold and 4-fold weaker than S454P or ΔNLS1 respectively (Fig. 3B), suggesting that NLS1 and the R3H-like motif cooperatively bind the same RNA. R507G and L398S mutations, which are outside of the R3H-like motif, did not compromise Tail capacity to bind RNA (Fig. 3B).

To determine if Tail binds RNA also in the context of the full length Xrn1, we employed an Electrophoretic Mobility Shift Assay (EMSA) and found that a single mutation in Tail RNA-binding domain, S454P, was sufficient to affect RNA binding of full-length Xrn1(Fig. 3C). These results suggest that the Tail RNA-binding motif (R3H-like) is a major component of the RNA-binding activity of Xrn1. To examine whether the 5’ end of the Tail-bound RNA binds the active site, we introduced the R101G mutation that was demonstrated to inactivate the interaction of the R101 with the 5’phosphate of the decaying RNA, thus weakening the interaction of the active site with the decaying RNA^3, 27, 28^. This mutation compromised the gel shift signal (Fig. 3C), indicating that the active site contributes to Xrn1 RNA binding capacity detected by our EMSA assay. Importantly, whereas a single mutation in R101G or S454P decreased RNA-binding by 60 or 65%, respectively, the R101G and S454P double mutant lost its RNA-binding capacity almost completely (Fig. 3C). This synergism suggests that the R101 and S454 bind the same RNA cooperatively. Being a major exonuclease, Xrn1 must bind RNA directly, at least in its active site. Xrn1 also binds directly full length RNA (Sohrabi-jahromi et al., 2019and our unpublished results). Other RNA-binding motifs and binding features are unknown. As discussed above, we found that Tail binds RNA in conjunction with R101 of the active site. To further examine whether Tail is involved in the major RNA-binding activity of Xrn1, we determined the repertoire of RNAs that Tail binds *in vivo*, using the CRAC technique, and compared it to that of the Xrn1^WT^. We found a significant correlation between the RPKM values of mRNA bound to Xrn1 and those that bound Tail (Fig. 3D), indicating that Xrn1 and Tail bind similar repertoire of RNAs. Taken together, the linkage between the active site and S454, and our observation that the tail binds the same repertoire of RNA as Xrn1 suggest that Tail provides a major RNA-binding site for the substrate RNA whose 5’ end is in the active site – the decaying RNA.

### The importin Kap120 is involved in nuclear import of decay-factors

For identifying Xrn1 importins, we scanned all known karyopherins by co-IP, comparing Xrn1 WT and Xrn1^ΔNLS1/2^, and singled out Kap120 (results not shown). Kap120 imports Rpf1 and Swi6 by binding their monopartite NLSs ^30^. To examine whether Kap120-Xrn1 interaction is direct, we affinity purified recombinant 6xHis-tagged Kap120 from *E. coli*, and FLAG-tagged Xrn1, or its mutant derivatives, from yeast extract (the purity of Xrn1 can be evaluated in Fig. 3C, middle panel). We reacted the two proteins *in vitro* and affinity purified the resulting complex and found that Xrn1 and Kap120 can interact *in vitro* and that Xrn1^ΔNLS1/2^-Kap120 interaction was 4.5-fold weaker than Xrn1^WT^-Kap120 (Fig. 4A), indicating that either NLS1 or NLS2 or both NLSs contribute to the *in vitro* interaction. The converse IP experiment led us to the same conclusion (Fig. S3A). Kap120 bound Xrn1^ΔNLS2^ better than Xrn1^ΔNLS1^ (Fig. 4B, designated in the figures as “ΔNLS1” and “ΔNLS2” respectively), indicating that, *in vitro*, NLS1 binds Kap120 better than NLS2. Kap120 also binds Tail, outside the context of Xrn1, in an NLS1-dependent manner (Fig. S3B). Co-IP experiment with yeast extract demonstrates that NLS1/2-mediated Kap120-Xrn1 interaction occur also *in vivo* (Fig. 4C). Using a similar CO-IP experiment, we found that Dhh1 and Pat1, but not Ccr4, also interact with Kap120 *in vivo* and, interestingly, this interaction was also ∼5-fold reduced in the Xrn1^ΔNLS1/2^ (Fig. 4D and results not shown).

**Fig.4.**
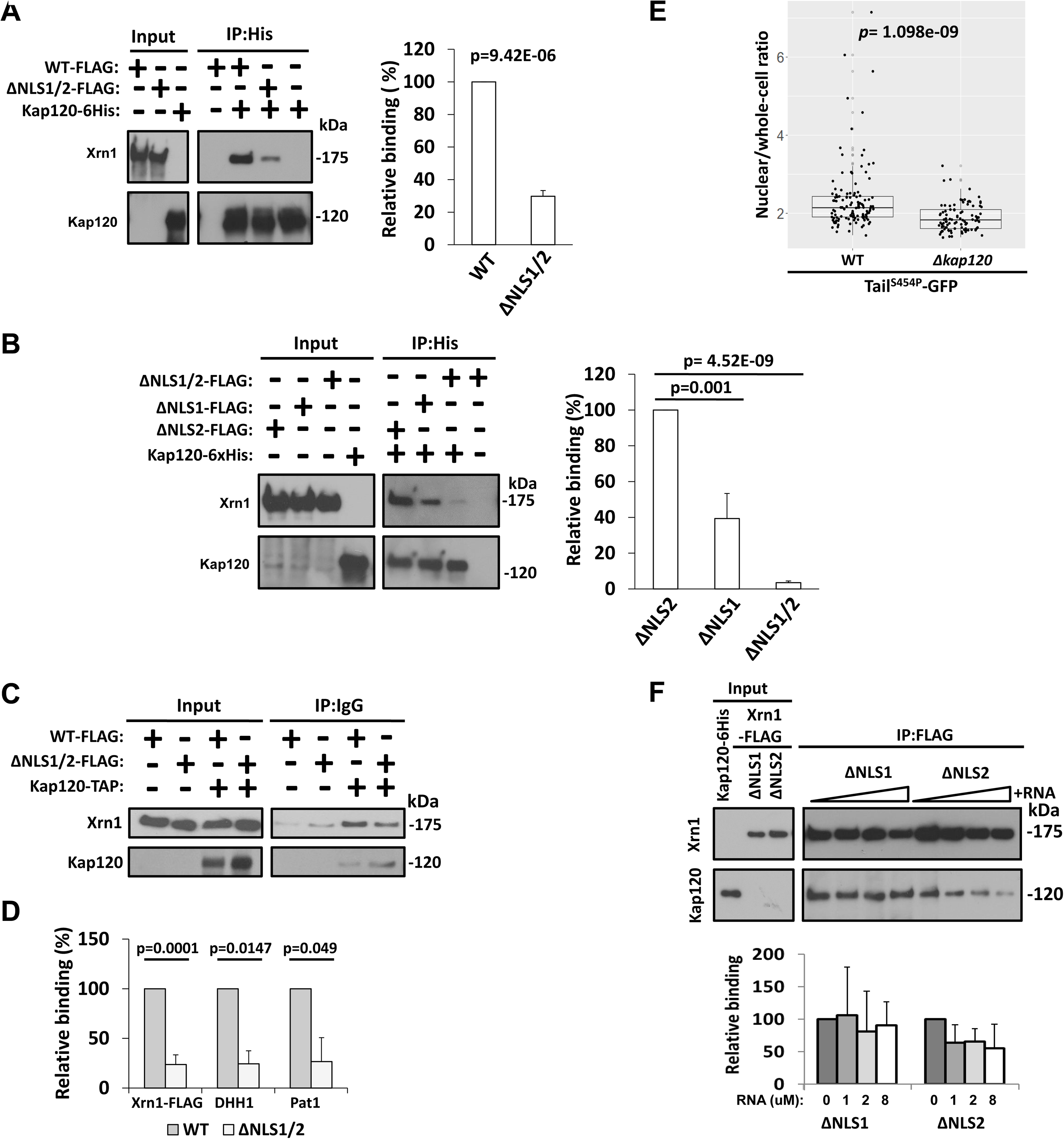
Kap120 recognizes Xrn1 NLSs. (**A**) *Interaction of Kap120 with Xrn1 is mediated by its NLSs*. Affinity purified FLAG-tagged Xrn1 or its mutant derivative, Xrn1^ΔNLS1/2^, and recombinant Kap120-6xHis that had been purified with Ni-NTA column were mixed together and co-IPed with Ni-NTA column, followed by western blotting, using antibodies against the indicated proteins. Western signal of 3 independent replicates were imaged using ImageQuant and quantified with TotalLab and plotted. Xrn1-FLAG intensity was normalized to Kap120 intensity, defining Xrn1^WT^/Kap120 as 100%. *p*-value was calculated using Student’s unpaired T-test. (**B**) *Kap120 binds both NLSs of Xrn1*. Experiment shown in A was repeated, in 3 replicates, except that Xrn1 molecules with mutations in a single NLS, as indicated, were examined. Right panel shows the quantification, performed as in A. Xrn1-FLAG intensity was normalized to Kap120 intensity, defining arbitrarily Xrn1^ΔNLS1/2^/Kap120 as 100%. (**C**) *Xrn1 interaction with Kap120 is mediated by NLSs in optimally proliferating yeast cells*. TAP-tagged Kap120 extracted from strains expressing either FLAG-tagged Xrn1,or its ΔNLS1/2 derivative, was affinity purified with IgG-sepharoseres in. The co-IPed proteins were subjected to Western blotting using the antibodies against the indicated proteins. Quantification is shown in D. (**D**) *NLS1/2 are required for efficient co-IP with Kap120 of Xrn1 and other DFs*. The membrane shown in C and two more membranes of additional replicates were also reacted with anti-Pat1 and anti-Dhh1 Abs. For each of the indicated proteins, its immunoblot western signal was subtracted with the signal of the no-tag control; the signal was normalized to that of Kap120.The normalized WT signal was defined as 100% and the normalized mutant signal was compared to the WT. *p*-value was calculated using Student’s unpaired t-test. (**E**) *Kap120 is used for efficient import of Tail^S454P^-GFP*. Optimally proliferating WT or Δ*kap120* cells expressing Tail^S454P^-GFPand *NUP49*-mCherry (to mark the nucleus) were inspected microscopically. The nuclear/whole-cell ratio of the fluorescent signal was determined by ImageJ, as described in Methods. The mean ratio (nuclear intensity/Mean whole-cell intensity) is represented in a jittered box-plot. The *p*-value was calculated using Wilcoxon rank sum test. (**F**) *RNA blocks Xrn1-Kap120 interaction*. Interaction between purified Kap120-6xHis and FLAG-tagged Xrn1 carrying mutations in either NLS1 or NLS2, as indicated, was determine as in A, except that increasing amounts of 40 b RNA and EDTA (to inactivate Xrn1) were included. Xrn1-Kap120 complexes were affinity purified by anti-FLAG antibodies and analysed and quantified as in A. Final RNA concentrations are indicated underneath the histogram.

Finally, we took advantage of the constitutive nuclear localization of Tail^S454P^-GFP (Fig. 2C) and deleted *KAP120* to examine its role in Tail^S454P^-GFP nuclear localization. Deletion of *KAP120* changed significantly the cellular distribution of Tail^S454P^-GFP (Fig. 4E), indicating that Kap120 plays a key role in Tail import. Yet, it could not abolish import, suggesting that, in the absence of Kap120, there is/are other karyopherin(s) that is/are capable of importingXrn1 – albeit less efficiently than Kap120.

Since NLS1 binds RNA (Fig. 3B, ΔNLS1), we determined the effect of RNA on the interaction of Kap120 with Xrn1^ΔNLS2^, carrying only NLS1, and found that it weakened the interaction (Fig. 4F, ΔNLS2 lanes). The RNA had little or no effect on the interaction of Kap120 with Xrn1^ΔNLS1^ (Fig. 4F, ΔNLS1 lanes), suggesting that NLS2 does not bind RNA in a manner that disturbs the interaction with Kap120. To corroborate the impact of RNA on NLS1, we focused on the Tail domain that contains NLS1. Consistently, RNA could outcompete Kap120-Tail interaction in a dose dependent manner (Fig. S3C). Collectively, Kap120 binds the two NLSs of Xrn1. RNA can outcompete the interaction of Kap120 with NLS1 but not with NLS2.

### Disruption of Xrn1 NLSs affects both mRNA synthesis and decay rates of ∼50% of the mRNAs

The mutations in NLS1/2 provide a powerful tool to study the possible linkage between mRNA degradation, import and transcription. Importantly, our mutations in Xrn1 NLSs neither change Xrn1 protein level (Fig. S4A) nor the overall growth-rate (Fig. S4B), and they do not affect Xrn1 enzymatic activity (see later).

As expected, blocking import by mutating both NLS1 and NLS2 (using the *xrn1*^ΔNLS1/2^ strain) compromised transcription rates (TRs) of >3000 genes (Fig. 5A, Table S3). *xrn1*^ΔNLS1^ strain was similarly defective in TR (Fig. 5A), albeit more modestly than the defect of *xrn1*^ΔNLS1/2^ strain.

**Fig.5.**
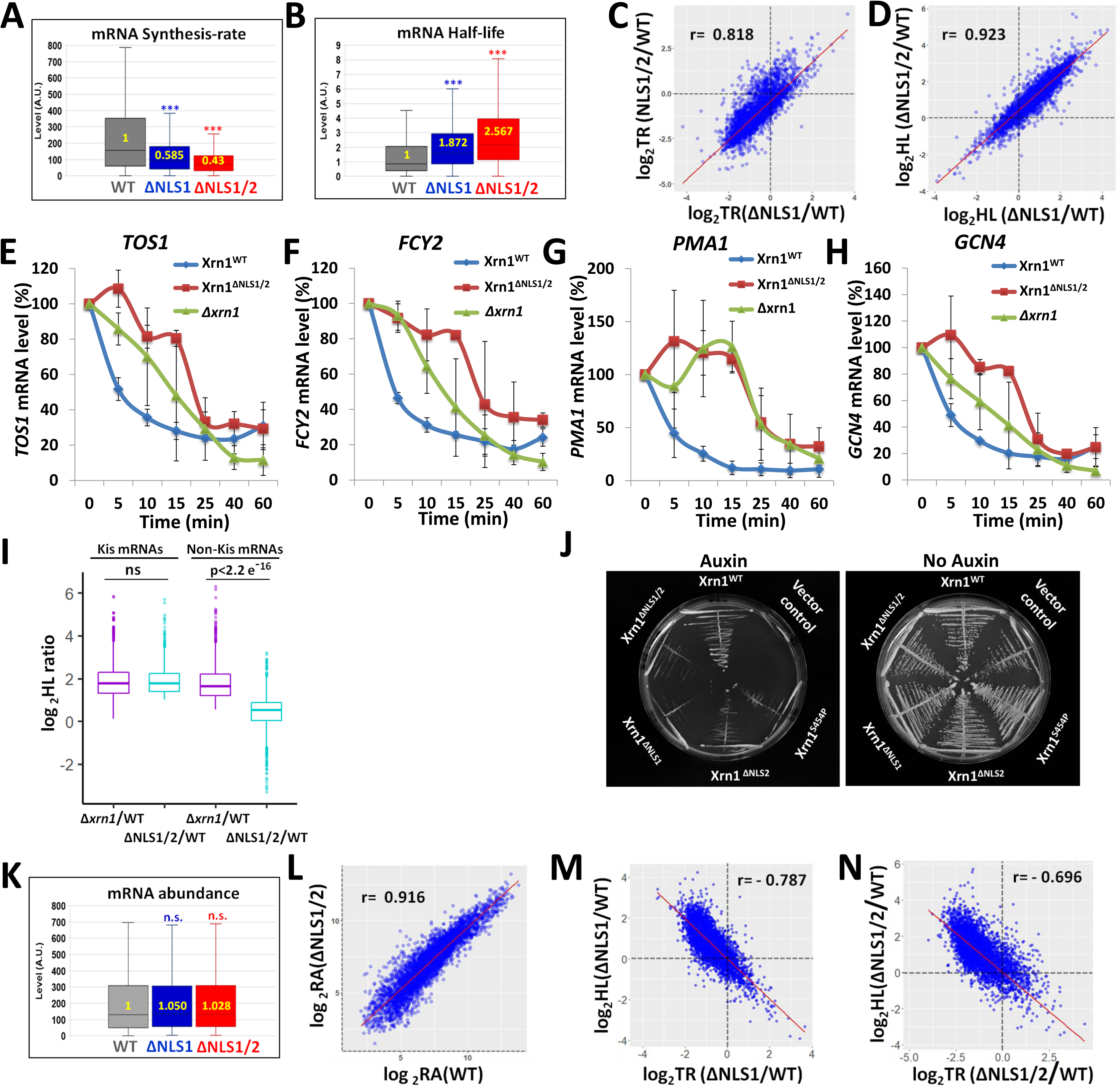
Blocking Xrn1 import leads to inefficient synthesis and decay of a large portion of the mRNAs, without affecting mRNA levels. Genomic Run-On (GRO) analysis was performed, in 3 replicates, as described in Method. Box and whisker plot of the median levels (in arbitrary units) of (**A**) *transcription rate (TR)*; (**B**) *half-lives (HLs)*. p-values were obtained by Wilcoxon rank sum test, ***: p<0.0001. (**C**) *Defect in TR of xrn1*^Δ^*^NLS1^ (expressed as a ratio between TR of the mutant and TR of WT) is correlated with that of xrn1*^Δ^*^NLS1/2^*. Scatter-plot of log_2_ mutant/WT ratio. Each spot represents an average of 3 replicates. Pearson correlation coefficient (r) and p value are indicated. (**D**) *Defects of mRNA decay (expressed as in C) of xrn1*^Δ^*^NLS1^ is correlated with that of xrn1*^Δ^*^NLS1/2^.* (**E-H**) *The effect of mutating NLSs on decay rates of specific mRNAs, determined by Northern blot hybridization*. Decay assay of the indicated mRNAs was performed as described in Methods and published previously. Shown are mRNA levels, quantified by PhosphorImager and normalized to *SCR1* mRNA, as a function of time post-transcription arrest. Results represent averages of 2 replicates (±S.D.) (**I**) *Effect of xrn1*^Δ^*^NLS1/2^ on Kis mRNAs HLs is comparable to that of XRN1 deletion, whereas it has little effect on HLs of non-Kis mRNAs*. Box and whisker plot of the log_2_HLs ratios between mutant and WT, as indicated below the boxes, for Kis and non-Kis mRNAs. Each box is based on 3 replicates. P-values were obtained by Wilcoxon rank sum test comparing WT and *xrn1^Δ^*^NLS1/2^ mutant. Ns – Not significant. Note that the two groups of mRNAs are equally affected by *XRN1* deletion. (**J**) *Proliferation of cells carrying mutations in Xrn1 NLSs is dependent on Ski2*. Synthetic sickness assay of NLSs mutants with *ski2Δ. ski2Δ, xrn1*-AID strains carrying either vector only, or a plasmid that expresses WT or the indicated mutant *XRN1* were streaked on a selective plate (to maintain the presence of the plasmid) in absence (right panel) or in presence of Auxin (that induced Xrn1-AID degradation). Photos were taken after 2 days at 30°C. (**K**) m<u>R</u>NA <u>A</u>bundance (RA) (steady state mRNA level) shown as in A. *p*-values were obtained by Wilcoxon rank sum test and were found to be not significant (n.s.). Median values in arbitrary unit (A.U.) are indicated inside the plots. (**L**) *RAs in xrn1^ΔNLS1/2^are correlated with those in WT*. Scatter-plot of log_2_ values of mutant and WT. Each spot represents an average of 3 replicates. Pearson correlation coefficient (r) is indicated. (**M-N**) *The effects of NLS1 (M) or NLS1/2 (N) disruption on transcription rates (TR) are correlated with their effects on HLs*. Scatter-plot of log_2_ mutant/WT ratio. Each spot represents an average of 3 replicates. Pearson correlation coefficient (r) is indicated.

Surprisingly, not only TRs were altered in the mutants, but also mRNA half-lives (HLs). More than 2600 mRNAs became ≥2-fold more stable in *xrn1*^ΔNLS1/2^ mutant compared to WT (Table S4). The median mRNA HLs in *xrn1*^ΔNLS1/2^ was 2.5-fold higher than that in the WT strain (Fig. 5B). We observed highly significant overlap between the mRNAs whose both synthesis and decay were ≥ 2-fold affected by blocking Xrn1 import (Fig. S4C). We named these 2401 overlapping mRNAs “Kem1 import sensitive” (“Kis”) mRNAs (Kem1 is an alias of Xrn1). Evidently, import of Xrn1 is required for both mRNA synthesis and decay. Thus, although blocking import seemed to permit more chance for the DFs to act (more Xrn1 molecules in the cytoplasm), they, in fact, acted as if there was less Xrn1 in the cytoplasm. This was the case for the Kis mRNAs. For the rest of the mRNAs, the “non-Kis” mRNAs, which are also degraded by Xrn1 (Xrn1 degrades most mRNAs^3, 8^), degradation was efficient (Table S5).

The effect of ΔNLS1 on TR and HL was correlated with the effect of ΔNLS1/2 on these parameters (Fig. 5C-D, respectively), consistent with the similar inhibition of Xrn1 import caused by ΔNLS1 and by ΔNLS2 (Fig. 1C). This strong correlation also suggests that disruption of NLS2 does not change the affected target genes.

A more direct experiment that determines HLs was performed by blocking transcription and monitoring levels of specific mRNAs following the transcription arrest. A number of mRNAs became more stable in *xrn1*^ΔNLS1/2^ (Fig. 5E-H) as well as in strains carrying a single NLS disruption (Fig. S4D-G). Collectively, this lack of a clear difference between ΔNLS1 and ΔNLS2, and the correlation between the strains (Fig. 5C-D), as well as the similar effect of ΔNLS1 and ΔNLS2 on Xrn1 import (Fig. 1C), suggest that the two NLSs function cooperatively in Xrn1 import and disruption of any one of them inhibits import (Fig. 1C) and affects the same repertoire of target genes.

These results prompted us to compare the effect on HLs of *xrn1*^ΔNLS1/2^ with that of *XRN1* deletion. Strikingly, we found no difference in HLs of Kis mRNAs by comparing *xrn1*^ΔNLS1/2^ with *Δxrn1* (Fig. 5I, “Kis mRNA”). In contrast, HLs of the non-Kis mRNAs were affected by *XRN1* deletion (as expected) but very little affected by blocking Xrn1 import (Fig. 5I, “non-Kis mRNA”). Thus, for Kis mRNAs, preventing Xrn1 import is analogous to a complete Xrn1 deletion, whereas non-Kis mRNAs are targeted for degradation by Xrn1^ΔNLS1/2^ almost as WT Xrn1. The two groups can be distinguished by the Gene Ontology (GO) terms of their products (Fig. S4I), most of which are related to ribosome biogenesis and composition, and the length of their open reading frame (ORF) (Fig. S4J). The efficient decay of the latter group in *xrn1*^ΔNLS1/2^ cells indicates that the enzymatic activity of Xrn1^ΔNLS1/2^ is not affected by the mutations in the NLSs. Note that the average HLs of the Kis and non-Kis mRNAs in *xrn1*Δ cells is comparable (Fig. 5I), indicating that the decay of both groups similarly requires Xrn1; the distinction between them is apparent only upon blocking Xrn1 shuttling. We also found no correlation between the effect that ΔNLS1/2 had on HL-ratio and the actual HL values in WT cells (Fig. S4H).

Cells require mRNA degradation activity for viability and cannot survive in the absence of the two major RNA exonucleases - Xrn1 or the exosome ^31, 32^. We introduced the auxin-induced degron (AID) tag ^33^ into *XRN1* (at its natural chromosomal locus) in cells lacking the exosome subunit *SKI2*. We used this system to verify the role of Xrn1 NLSs in mRNA decay by introducing plasmids expressing either XRN1 or its NLS mutant derivatives or a vector control into this *Δski2, XRN1*-AID strain. In the absence of auxin, these *Δski2, XRN1*-AID cells grew normally (Fig. 5J, “No auxin”). Plating these cells on auxin-containing plates stimulated Xrn1-AID degradation whose sole source of Xrn1 was the plasmid borne Xrn1. We found that a mutant form with disruption of a single NLS, or disruption of both NLSs, became sick when plated on Auxin (Fig. 5J, “Auxin”). The synthetic sickness of NLS mutants with *Δski2* is consistent with a defect of the NLSs mutants in mRNA decay. We suspect that, incase Xrn1 cannot import and Kis mRNAs cannot be degraded by Xrn1 (Fig. 5I), the exosome becomes the main degradation machinery of Kis mRNAs.

### Under optimal proliferation conditions, Xrn1 import does not affect mRNA steady state levels

Intriguingly, the average steady state (SS) level (Fig. 5K), and the SS level of most mRNAs (Fig. 5L) remained constant in all the studied strains. Thus, under optimal environmental conditions, shuttling of Xrn1 affects the dynamics of mRNA biogenesis and turnover without changing mRNA levels. A significant correlation was found between the effect of blocking Xrn1 import, using *xrn1*^ΔNLS1^ or *xrn1*^ΔNLS1/2^, on TR and its effect on HL (Fig. 5M-N). Thus, the more mRNA synthesis rate is dependent on Xrn1 import, the more its decay rate is affected. Collectively, the capacity of Xrn1 to degrade mRNAs is related to its capacity to stimulate their synthesis.

### Xrn1 nuclear import regulates exit from quiescence and proliferation under fluctuating (oscillating) temperature

We have examined several environmental conditions (e.g., high salt, high osmolarity, high pH, low pH, high temperature, low temperature, ethanol or glycerol as the sole carbon source) and found out that under many of which disruption of Xrn1 shuttling had little effect on the proliferation rate (in contrast with XRN1 deletion). However, when we plated cells that had experienced long-term starvation (≥7 days), we observed that *xrn1*^ΔNLS1/2^ colonies were smaller than WT (data not shown), raising the possibility that they exited starvation abnormally slowly. A characteristic feature of yeast cells that exit the starvation is increase in the proportion of budded cells. Indeed, budding proportion of WT cells increased during exit, exhibiting two budding phases due to cell synchronization. In contrast, budding of *xrn1*^ΔNLS1/2^ was delayed and unsynchronized (Fig. 6A). We also used Flow Cytometry to view the progression into the cell cycle. This assay showed that starved WT cells (“time 0”) consist of 75% cells arrested with 1 copy of their genome (1N, i.e.,G_0_ or G1) and virtually no S-phase cells, where as *xrn1*^ΔNLS1/2^ cells consist of 70% cells in G_0_ or G1 and ∼12% cells in S-phase. This suggests that the mutants did not arrest properly in G_0_ and some cells stopped dividing before DNA replication was completed, or started replication during starvation. More importantly, by 3.75 h and to a greater extent at 6 h post re-feeding, the WT cells started to replicate their genome and exiting stationary phase, whereas the mutant cells exhibited abnormally delayed signs of exiting starvation (Fig. 6B-D). In contrast, during optimal proliferation conditions when cells proliferated exponentially in rich medium, we observed no differences in DNA content of the two strains (Fig. S5A).

**Fig.6.**
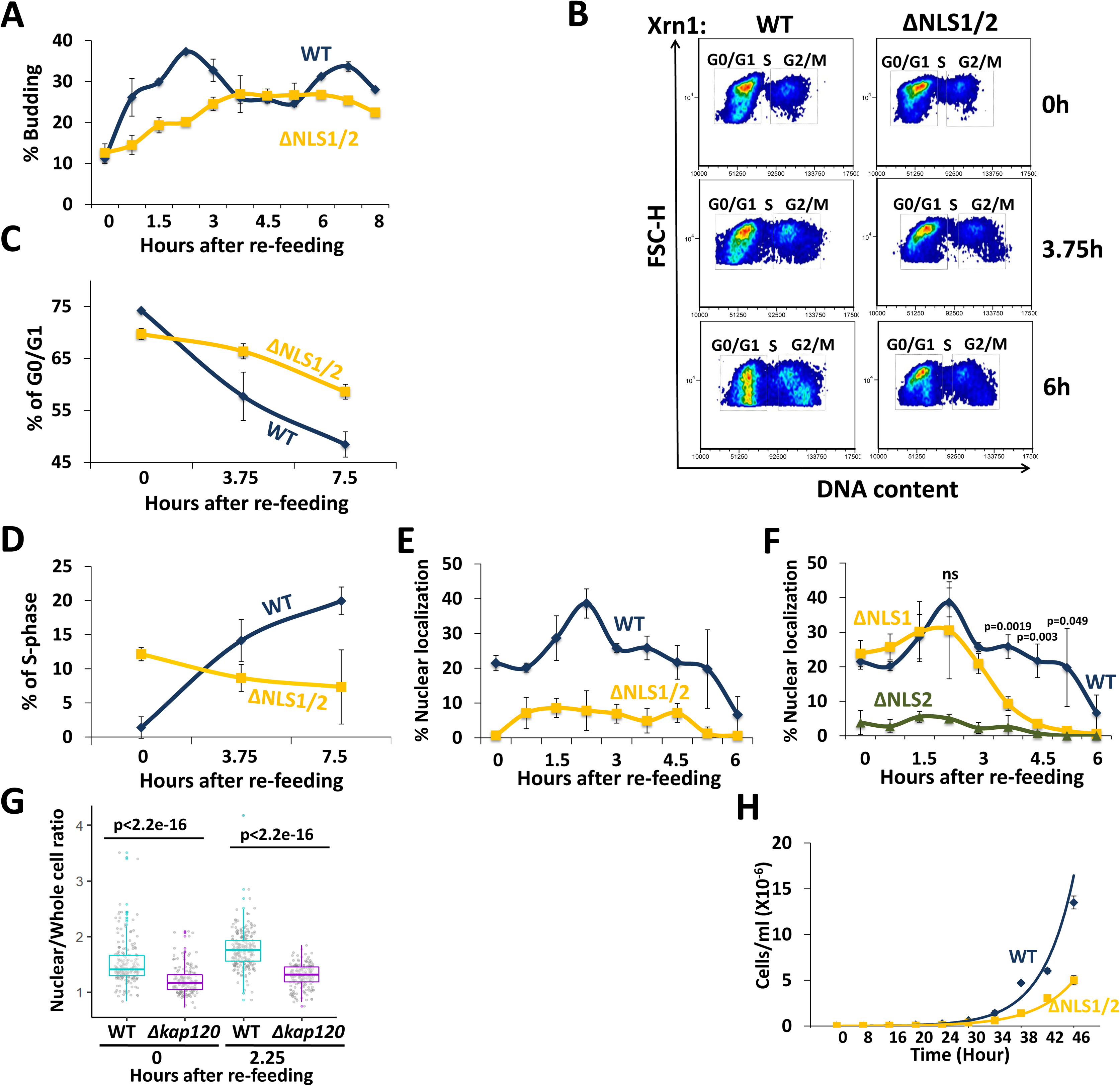
Xrn1ΔNLS1/2 cells exit starvation slowly and abnormally upon re-feeding, and are defective in proliferation at fluctuating temperatures. (**A**) *xrn1^ΔNLS1/2^mutant cells bud abnormally slowly during exit from long-term starvation to sated conditions*. WT and mutant cells were allowed to proliferate in YPD medium at 30°C until they both entered, simultaneously, into stationary phase (starvation). Seven days later, the cultures were diluted 10-fold in fresh YPD medium. Percentage of budded cells was plotted as a function of time post re-feeding (post dilution). (**B-D**) *xrn1^ΔNLS^^1/2^ mutant cells enter abnormally slowly into S-phase, as assayed by FACS analysis*. Cells were re-fed as in A. DNA content was determined by Flow cytometry (FACS), using CYTOX^TM^ Green, and plotted as a function of time post-re-feeding. Cells with 1N DNA content represent G_0_ or G1 cells. As these haploid cells exit starvation and enter S-phase, their DNA content gradually increases (**B**) and the % cells that enter S and G2 phases gradually increases (**C-D**). (**E**) *Xrn1^WT^-GFP, but not Xrn1^ΔNLS1/2^-GFP accumulates in the nucleus concomitantly with bud appearance*. Cells were inspected under a fluorescence microscope and a % cells with nuclear localization of GFP was plotted as a function of time following re-feeding. Error bars represent S.D. of 3 replicates. Note that, in this import experiment, no cycloheximide was added to avoid its effect on the actual process of exiting the starvation. Therefore, in this case (unlike our other experiments), we cannot differentiate between import of pre-existing Xrn1 or newly synthesized one. (**F**) *NLS2 is required for Xrn1 import during starvation and exit from starvation, whereas NLS1 is required only following 3 h post re-feeding*. Experiments of the four indicated strains shown in E and F were performed in parallel. Results were analysed as in E. The indicated *p*-values are the results of student’s t-tests, based on 3 replicates, between WT and ΔNLS1 strains at each indicated time-points (at 6 h, the difference was not significant). (**G**) *Kap120 is required for efficient import of Xrn1 during starvation and exit from starvation*. WT or Δkap120 cells, expressing XRN1-GFP and RPB7-RFP (to mark the nucleus), were inspected microscopically. After examining the starved cells (Time 0), the culture was re-fed as in E and examined microscopically at 2.25 h later. The nuclear/whole-cell ratio of the mean fluorescence intensity was determined by ImageJ for each individual cell, as described in Methods. P-value, based on 3 replicates, was calculated by Wilcoxon rank sum test. (**H**) *Xrn1 import is required for normal proliferation rate under fluctuating temperatures*. WT and *xrn1*^ΔNLS1/2^ cells were cultured in 100 μl of rich liquid medium (YPD) in many PCR tubes (each was used for measuring one time point), at initial 10^4^ cells/ml. Tubes were placed in a PCR machine. After 3 h acclimation at 30°C, the temperature began to cycle between 38°C for 150 min and 8 °C for 150 min. At the indicated time points, a tube was taken and cell density was determined by counting cells using haemocytometer. Both strains proliferated comparably at constant temperatures (see Fig. S5D-F).

These results provoked us to examine Xrn1 localization post-refeeding. We found that ∼20% of starved cells accumulated Xrn1 in the nucleus, as reported previously ^3^. During the first phase of exit from starvation (the kinetics is dependent on the time that cells experience the starvation), cells continued to accumulate nuclear Xrn1 and the proportion of nuclear localized cells gradually increased. Later, the nuclear localization gradually decreased until it reached the baseline that characterizes optimally proliferating cells (Fig. 6E “WT”, Fig. S5B, “Xrn1^WT^-GFP” panels). As expected, disruption of NLS1/2 resulted in little nuclear localization (Fig. 6E-F, Fig. S5B-C). Interestingly, NLS2 but not NLS1 was responsible for importing Xrn1 in response to re-feeding (Fig. 6F). Xrn1^ΔNLS1^-GFP localization paralleled that of Xrn1-GFP until 3h post re-feeding. Later, during entry into log phase, NLS1 became an important NLS (Fig. 6F, >3 h), as we observed in optimally proliferating cells (Fig. 1C). Thus, starvation induced import of Xrn1 is specifically mediated by NLS2. Moreover, during the transition from starvation to rich medium, there is an additional wave of import that is specifically dependent on NLS2 as well. Not only was Xrn1 imported during the exit from starvation, but also Pat1 was imported concurrently with Xrn1 in an NLS1/2-mediated fashion (Fig. S5C). Kap120 was involved in Xrn1 localization during starvation, as well as in the additional import observed during the first phase of exit from starvation that peaked at 2.25 h post-refeeding (Fig. 6G). Taken together, Xrn1 import is required for proper exit from stationary phase in response to re-feeding. Whereas, during optimal proliferation, both NLS1 and NLS2 are required for efficient import, during long-term starvation and exit from it, NLS2 seems to play the major role.

The impact of Xrn1 import on the cell responses to changes in nutrient availability, which characterizes the transition from starvation to sated conditions, encouraged us to examine cell proliferation rate under conditions that simulate continuous environmental changes. For simplicity we chose fluctuating temperatures. Exponentially proliferating cells were cultured in a thermal cycler, enabling us to oscillate the temperature between heat shock (38 °C for 150 min) and cold shock (8 °C for 150 min). Each of these temperatures, although extreme for yeast growth, permitted cell proliferation with no difference between the strains when used continuously (Fig. S5D-F, and results not shown). However, when the cultures were fluctuated between these temperatures, *xrn1*^ΔNLS1/2^ grew slower than WT cells (Fig. 6H). Collectively, we find that the shuttling feature of Xrn1 is not required for proliferation during continuous conditions with an unchanging environment, however harsh; it is required mainly, if not exclusively, for proper and rapid responses to environmental changes.

## Discussion

The eukaryotic cell contains two major cellular domains, nucleus, and cytoplasm, the linkage of which involves the transport of various molecules. The importance of this linkage is exemplified by the mechanisms that control mRNA levels, which are mediated by a balance between RNA synthesis and decay. We hypothesized that proper linkage between these two opposing processes involves shuttling of factors that promote mRNA synthesis and decay: transcription factors and mRNA decay factors. As the mRNA decay complex mediates both processes, we surmised that its shuttling is instrumental for the linkage between the nuclear mRNA synthesis and the cytoplasmic decay. Here we uncover a nucleocytoplasmic shuttling mechanism of Xrn1 and probably some of its associated factors that are regulated, in part, by the bound RNA, which in turn, regulates mRNA synthesis and decay.

### Under optimal conditions, NLS1 and NLS2 seem to function cooperatively

Bioinformatic analyses revealed two NLSs in Xrn1, but none in a number of other known DFs that we examined. We verified these NLS sequences are necessary for nuclear shuttling and showed that these NLSs are recognized by Kap120. Although, *in vitro*, Kap120 binds NLS1 better than NLS2 (Fig. 4B), disruption of any one of them compromised Xrn1 import substantially (Fig. 1C) and resulted in defective mRNA decay rate (Fig. S4D-G). Consistently, the effect of disruption of a single NLS on TRs and HLs is correlated with that of disruption of both NLSs (Fig. 5C-D). These data suggest that the two NLSs are cooperatively required for Xrn1 import and disruption of any one of them inhibits import. The underlying mechanism of cooperativity remains to be determined. Are two NLSs needed to cover Xrn1 from its different ends to permit efficient transport through the nuclear pore, as was proposed in other cases of large proteins ^34^? In contrast to optimal proliferation conditions, upon starvation the two NLSs seem to acquire more distinct functions. Disruption of NLS2, but not NLS1, inhibited Xrn1 import. This was observed both during long-term starvation (Fig. 6F time 0) and during exit from starvation, upon re-feeding (Fig. 6F), conditions of which induced an additional wave of Xrn1 import (Fig. 6A, F-G). We therefore suspect that the proposed cooperation between NLS1 and NLS2 does not operate under all environmental conditions and may be subjected to regulation.

### Nuclear Xrn1 is required for cytoplasmic decay of ∼2400 “Kis” mRNAs

The vast majority of yeast mRNAs are degraded mainly in the cytoplasm by an Xrn1-dependent pathway ^3, 5, 8^. Prior to this work, we hypothesized that preventing Xrn1 import would decrease mRNA synthesis of target genes and increase decay of these gene transcripts, because more Xrn1 would be available in the cytoplasm for mRNA decay. Our results proved otherwise. Specifically, NLS1/2 mutant cells exhibited slow mRNA decay rate. Strikingly, the decay rates of the client mRNAs, named Kis mRNAs, were comparable to those in Δxrn1cells (Fig. 5I). We therefore propose that, for degradation of Kis mRNAs by Xrn1, Xrn1 needs to enter the nucleus. Because the impact of *xrn1*^ΔNLS1/2^ on HLs is comparable to that of *Δxrn1* (Fig. 5I), we surmise that if Xrn1 is not imported, Kis mRNA are degraded only by an Xrn1-independent pathway, e.g., the exosome (Fig. 7B, left pathway). Indeed, *xrn1*^ΔNLS1/2^ cells become sick upon deletion of *SKI2* (encoding an exosome component) (Fig. 5J).

**Fig. 7.**
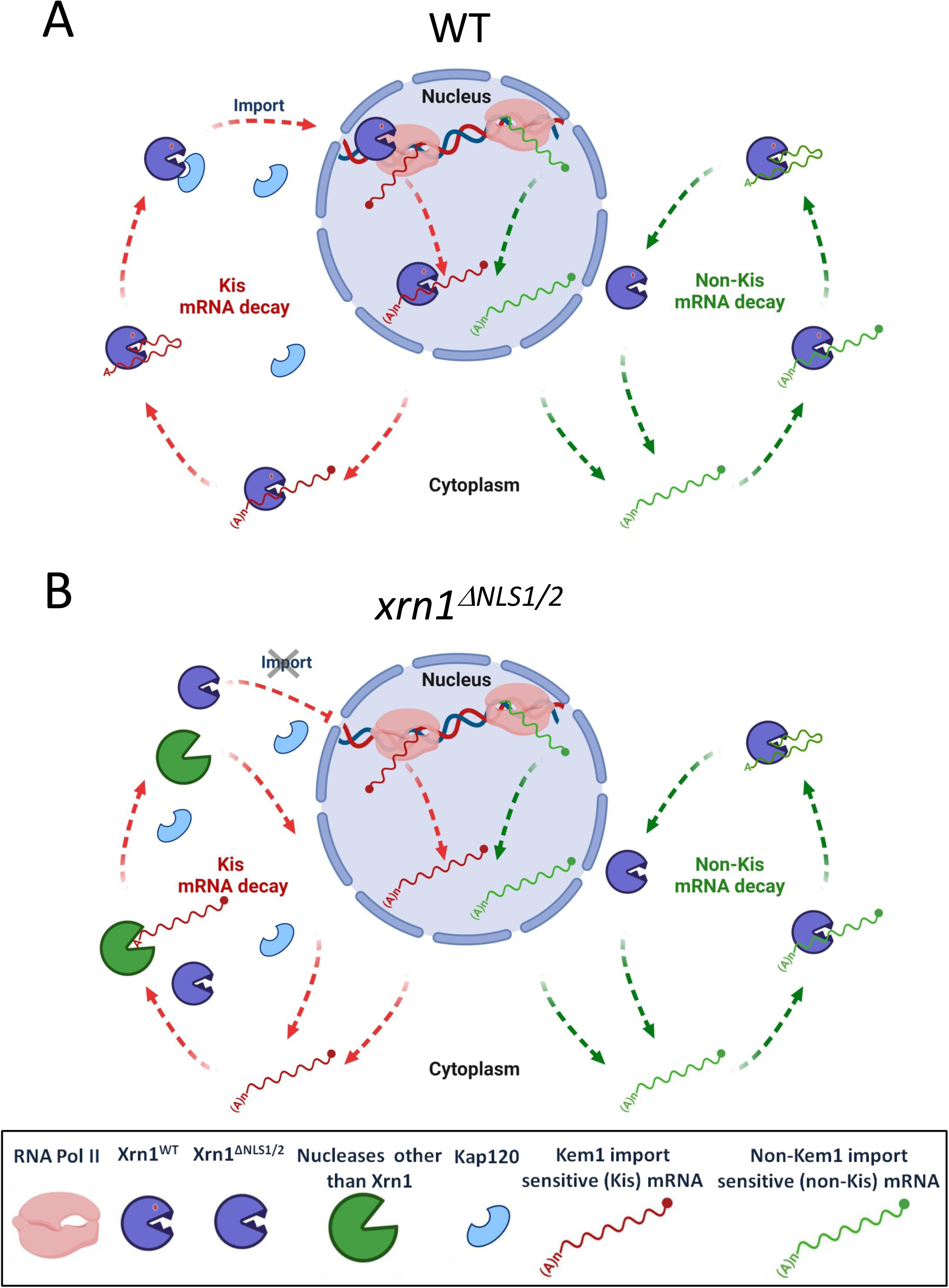
A proposed model for two Xrn1-mediated mRNA decay pathways, one of which begins in the nucleus. (**A**) *WT cells*. Left pathway (red arrows) illustrates the Kis mRNAs pathway, which begins in the nucleus. Xrn1 binds promoters and stimulates transcription ^1–3^. During transcription, it binds the emerging transcript ^68^, most probably at its 3’ end^29, 69^ (also our unpublished results). This RNA is protected by the cap, yet Xrn1 is immediately available upon decapping, or after endonucleolytic RNA cleavage. In this model, we hypothesize that Xrn1 does not dissociate from binding the 3’ end. Thus, upon decapping, Xrn1 binds both the 5’P and the 3’ RNA end (until advance stage of the RNA degradation). This hypothesis remains to be examined experimentally. Following RNA decay, NLS1 is exposed which binds Kap120, and Xrn1 is imported to begin a new cycle. Right pathway (green arrows): the decay of non-Kis mRNAs is, by definition, insensitive to Xrn1 shuttling. It is confined to the cytoplasm and follows the standard model of mRNA decay ^8, 9^. We do not know whether the Xrn1 molecule that degrades these mRNAs are imported to the nucleus. (**B**) *xrn1*^Δ^*^NLS1/2^cells*. Because the impact of *xrn1*^ΔNLS1/2^ on HLs is comparable to that of *Δxrn1* (Fig. 5I), Kis mRNAs are degraded mainly by Xrn1-independent pathway. We note that blocking import increases the number Xrn1 molecules in the cytoplasm (represented by more Xrn1 icons) and seems to permit more chance for Xrn1 to act in the cytoplasm; however, this surplus Xrn1 does not seem to enhance degradation (depicted as “idle” Xrn1 icons in the red pathway). See text for more details.

Degradation of the “non-Kis mRNAs” is little dependent on Xrn1 shuttling (Fig. 5I) suggesting that recruitment of Xrn1 to these mRNAs occurs in the cytoplasm. Normal decay of the latter mRNAs indicates that Xrn1^ΔNLS1/2^ is enzymatically functional. This conclusion is supported by the WT growth rate of *xrn1^Δ^*^NLS1/2^ cells under many different constant environmental conditions (but not during fluctuating environments – see below).

Kis and non-Kis mRNAs can be distinguished by the effect that Xrn1 import has on (i) their TR (this was one of the criteria of their definition), (ii) the Gene Ontology (GO) terms of their products (Fig. S4I); most of the Kis GO terms are related to ribosome biogenesis and composition, whereas the GO terms of non-Kis genes were different and exhibited little connectivity, and (iii) the length of their open reading frame (ORF) (Fig. S4J).The efficient decay of the non-Kis mRNAs in *xrn1*^ΔNLS1/2^ cells indicates that the enzymatic activity of Xrn1^ΔNLS1/2^ is not affected by the mutations in the NLSs. It is worth noting that there was no correlation between the effect that ΔNLS1/2 had on HLs (ratio) and the HL values in WT (Fig. S4H). Thus, the distinction between Kis and non-Kis mRNAs is unrelated to their actual HLs.

### A linkage between RNA decay and Xrn1 import

We also uncovered an inverse linkage between mRNA degradation and transcription, whereby binding of RNA to NLS1 serves as a mechanism to regulate Xrn1 import. More specifically, we provide evidence that RNA (containing 5’phosphate), which binds both the active site and the R3H-like motif in a cooperative manner (Fig. 3C, compare lanes “R101G” and “S454P”, with the lane “R101G, S454P”), also binds NLS1 (Fig. 3B, see lane “ΔNLS1”). Moreover, RNA inhibits the *in vitro* Kap120-Xrn1 interaction (Fig. 4F). This establishes a retrograde linkage between mRNA decay and Xrn1 import and transcription. As the two NLSs are required for import, we propose that as long as the RNA is covering NLS1, import is repressed; only following RNA degradation, NLS1 is exposed and Xrn1 is imported to the nucleus. Interestingly, blocking Xrn1 import, which increases the Xrn1 concentration in the cytoplasm, did not contribute much to degradation of non-Kis mRNAs because only a minor population of mRNAs (∼100) become >2-fold unstable in the NLS1/2 mutant (see Table S4).

### Xrn1 shuttling plays a role in the responses to environmental changes

During constant environmental conditions, Xrn1 shuttling plays little role in mRNA buffering and has no effect on proliferation rates. However, Xrn1 shuttling plays a role during transitions from one condition to another. Two examples are shown here. The first is a transition from starvation (stationary phase) to sated conditions. Indeed, in such cases *xrn1*^ΔNLS1/2^ cells exit more slowly and abnormally from starvation (Fig. 6A-D). The second example is fluctuating temperatures. Fluctuating environments such as changes in ambient temperature or nutrient availability represent a fundamental challenge to living organisms, including yeast. Under these ever-changing conditions, that better represent the real yeast habitat, the proliferation rate of *xrn1*^ΔNLS1/2^ cells is abnormally slow. In contrast, the mutant cells can proliferate normally at any temperature that we tested, provided that it does not change. In response to various stresses, the level of mRNAs transiently increases, resulting in a “peak behaviour” ^35–39^. This is important for properly coping with the stress (Shalem et al., 2011 and references therein). It was previously demonstrated mathematically that, given a constant TR, the kinetics of the peak behaviour in response to the environment is solely dependent on mRNA decay rate; a faster rate results in a faster response, i.e., the time it takes for mRNA levels to reach their highest value is directly affected by the decay rate ^40, 41^. In case both mRNA synthesis and decay rates increase, the kinetics of mRNA levels changes are even faster ^40^. Therefore, it seems that the effect of Xrn1 import on enhancing both mRNA synthesis and decay in response to environmental changes is related to the kinetics of the response and not to the SS levels of these mRNAs after the cells adapt to the new environment. This is in line with the proposal that higher turnover allows faster response to the environment regardless of the actual SS mRNA levels ^40, 42^. We therefore propose that a main function of Xrn1 shuttling (and its mRNA decay associated proteins) under changing conditions is to permit rapid acquisition of new SS levels. However, once the new levels are obtained, Xrn1 shuttling is dispensable for subsequent proliferation rates. Yeast cells can obtain homeostasis in response to all kinds of changes, if enough time is provided ^43^. The cell seems to be a plastic system, but this feature is not enough for competing with other cells in the wild. The rate of obtaining this homeostasis is likely critical for competition. We hypothesize that proper and rapid shuttling of molecules between the two major compartments has been evolved, in part, to permit rapid response to the environment and perseverance in the face of constant environmental uncertainty.

## Methods

### Yeast Strains and Plasmids construction

Yeast strains, and plasmids are listed in Supplementary Tables 1 and 2 (Table S1 and S2). Xrn1 NLSs were mutated by homologous recombination with PCR amplified mutated fragments, primers carrying mutations were used to incorporate the mutations. For creating yeast strains, Xrn1 (or, gene of interest) is replaced by *CaURA3*. WT or mutated fragments was later introduced by replacing *CaURA3* by homologous recombination and selection on 5-FOA. In this way, mutations were surgically introduced without adding other sequences. All strains were verified by both PCR and Sanger sequencing.

### Yeast proliferation conditions, under normal and fluctuating temperatures, and during exit from starvation

Yeast cells were grown in synthetic complete (SC), Synthetic dropout (SD) or in YPD medium at 30°C unless otherwise indicated; for harvesting optimally proliferating cells, strains were grown for at least seven generations in log phase before harvesting. For nucleocytoplasmic shuttling assay, cells were grown at 24°C before shifting to 37°C for 2 hr. For nutrient starvation experiment, we employed previously described protocol^3^; briefly, exponentially growing cells were washed once with water and then resuspended in Synthetic media lacking both carbon source and amino acids. Cells were imaged after 1h with a fluorescence microscope. For exit from long-term starvation, cells were proliferated in SD medium at 30°C and allowed to enter stationary phase. Following entry, cells were incubated for 7 additional days. To re-feed the culture, cells were collected by centrifugation and were resuspended in fresh medium at 10 fold smaller cell density, followed by incubation at 30°C. Yeast proliferation was measured by counting the buds with a phase-contrast microscope. For fluctuating temperature assay, cells were grown in YPD over-night till mid log phase. Cells were then cultured in 100μl of liquid rich medium (YPD) in several PCR tubes, each tube was used for measuring one time point, at initial 10^4^ cells/ml. After 30 min acclimation at 30°C, the temperature cycled between 38 °C for 150 min and 8 °C for 150 min for indicated time.

### Synthetic sickness assay

Optimally proliferating cells that were taken from tiny colonies, developed during over-night incubation at 30°C on an SC plate, were streaked on plates with or without auxin and incubated for 2 days at 30°C prior to photography.

### Nucleocytoplasmic Shuttling Assay

Cells were allowed to proliferate at 24°C until 3-5 × 10^6^ cells/ml, rapidly shifted to 37°C and subsequently incubated for up to 2h. The translation inhibitor cycloheximide (CHX) (Sigma) was added to a final concentration of 300µM just prior to temperature shift. Pab1p-GFP, a shuttling protein^22^ was used to determine the efficiency of the assay and to serve as a nuclear marker^3, 22^. In all cases, Pab1p-GFP was present in the nuclei of at least 60% of the heat inactivated mutant cells after 2 hr.

### Identifying Tail mutations that led to constitutive nuclear localization

A PCR based random mutagenesis protocol^44–46^ was used to mutate Tail-GFP fragments. The mutants were introduced into a WT strain (yMC229) and analysis was performed by fluorescence microscopy followed by sequencing of nuclear-localized isolates.

### Fluorescence microscopy

Fluorescence microscopy was performed as described before^3, 47^. Image processing and quantitation when required was done by ImageJ^48, 49^. The nuclear/whole-cell ratio of the fluorescent signal was determined by ImageJ. The nuclear and cellular boundary (ROI) was defined by hand-drawn tool of ImageJ, followed by measuring the intensity

### Affinity purification of Kap120-HTP, Xrn1-FLAG and *in vitro* interaction assay between the purified proteins

Cells were grown in YPD media until 2-2.5 x 10^7^cells/ml and were harvested by filtration and flash frozen by plunging into liquid nitrogen. Frozen cells were lysed cryogenically via six cycles of pulverization (15 herz) using a mixer mill 400 (RETSCH). Grindates were later resuspended in appropriate lysis buffer supplemented with protease inhibitors, centrifuged (8000 g for 10 min) and protein amount was measured. Subsequently equal amounts of protein were subjected to IP. FLAG-IP was done using FLAG kit (Sigma, Cat.No: FLAGIPT1), according to manufacturer’s protocol. Briefly, protein samples were rotated overnight at 4°C with anti-FLAG M2 sepharose beads. Sepharose suspensions were centrifuged for 30 seconds at 1000 g, supernatant discarded and subsequently washed 4 times with 1X Wash Buffer. IP was eluted twice with 3X FLAG Elution Buffer and stored at −70°C until further use. Protein-A tagged proteins were affinity purified by incubating with IgG sepharose resin for 2 hr at 4°C. The resins were washed 4 times with IPP150 (10 mM Tris-Hcl pH 8.0,100 mM KAc, 0.1% NP-40, 2 mM MgAc,10 mM β-Mercaptoethanol, 1 mM PMSF), followed by washing once with TEV cleavage buffer (10mM Tris-HCl pH 8.0, 100 mM KAc, 0.1% NP-40, 1 mM MgAc,1 mM DTT) without the TEV. The elution was made with by incubating the beads in TEV buffer with TEV protease for 2h at room-temperature. Eluates were stored at −70°C until further use. His-tagged protein purification was done with Ni-NTA agarose resin (Qiagen) according to manufacturer’s protocol; briefly, overnight bacterial culture with Kap120-6xHis was grown for few hours. At 0.6 O.D., 0.1 mM IPTG was added. Four h later, cells were harvested and resuspend very gently with 10 ml ice cold 1x PBS, spinned 8000 rpm for 10 min at 4°C. The pellet was frozen at −80°C. To extract proteins, the frozen cell pellet was resuspended in ice cold 30 ml of lysis buffer (50mM Tris-HCl, pH 8.0, 250 mM NaCl, 0.5% Triton-X-100,10mM Imidazole + protease inhibitors). Cold cells were subjected to 6 pulses of sonication −10 sec each-with intervals of 30 sec in ice, spinned for 5min at 13000 rpm 4°C. Seven ml of unpacked Ni-NTA resin was washed with 2 x 7 ml lysis buffer and the supernatant was mixed with the resin for 1 hr in a rotator at 4°C. The mix was transferred to a column and beads were washed with 3x 10 ml Wash-buffer (50mM Tris-HCl, pH 8.0, 250 mM NaCl, 0.5% Triton-X-100, 20 mM Imidazole, pH was adjusted to 8.0 using NaOH). Protein was eluted twice with 1 ml elution buffer (50mM Tris-HCl, pH 8.0, 250 mM NaCl, 0.5% Triton-X-100, 250 mM Imidazole, pH was adjusted to 8.0 using NaOH).

For *in-vitro* interaction assays, purified His-tagged Kap120 was mixed with Xrn1-FLAG (WT or mutant) protein and incubated in a rotator for 1 h at 4°C. To pull down Xrn1-FLAG/Kap120-HISx6 complex, anti-FLAG coated beads or Ni-NTA agarose beads were used, and proteins were eluted with 2X LSB and heat. For experiments involving RNA, a commercially synthesized 40 nt long RNA (5’-AUGUACAGUCCGACGAACACGGAGUCGACCCGCAACGCGA-3’) was added to the mix, EDTA (2 mM final concentration) was added to prevent degradation of RNA by Xrn1. Similarly, GFP-tagged Tail was pulled down with GFP-Trap^®^ resin (ChromoTek) in a small column, washed 3 times with 1X Wash buffer. Purified Kap120-His was added to the columns and incubated in a rotator for 1 h at 4°C. The column was washed and eluted with 2X LSB at 95°C.

### Western blotting analysis

Proteins were run in a precast SDS-PAGE gel (BioRad), followed by an electro-transfer to a Nitrocellulose membrane (BioTrace^TM^ NT; PALL Life Sciences). The membrane was blocked for 1 hour with non-fat milk at room temperature or over-night at 4°C. It was then reacted with appropriate primary antibody O.N at 4°C, or 1 h at room temperature, washed and subsequently reacted with HRP conjugated secondary antibody. Band intensity was detected with the Western Lightning Plus-ECL (Perkin Elmer) according to the manufacturer’s instructions. The membrane is exposed to either X-ray film or photographed with ImageQuant^TM^ LAS4000 (GE Healthcare).

### RNA-Protein interaction assays

One liter of culture was harvested at 1 x10^7^ cells/ml and grinded cryogenically as described above. For UV crosslinking, the grindat was transferred to a glass petri-dish and mixed with powdered dry-ice, irradiated 3 times in an UV-crosslinker (Stratalinker) with 0.6 J/cm^2^. During intervals, more dry-ice powder was added to replenish evaporated dry-ice, keeping the cell powder frozen during the crosslinking. The grindate was then suspended in lysis buffer (6 mM Na_2_HPO_4_, 4 mM NaH_2_PO_4_.H_2_O, 1% NP-40, 100 mM Potassium Acetate, 2 mM Magnesium Acetate, 50 mM Sodium Fluoride, 0.1 mM Na_3_VO_4_, 10 mM β-mercaptoethanol, protease and phosphatases inhibitors, centrifuged at 20000g for 15 min at 4°C and supernatant was collected. Equal amount of proteins were mixed with IgG sepharose beads, incubated for 1 hr at 4°c. The samples were later transferred to the small empty columns (Bio-Rad) and washed 3 times with lysis buffer and once with TEV cleavage buffer. 1U of RNase-cocktail (RNaseA + T1, Thermo) was added to the beads for 5min at 37°C, reaction was stopped by adding 650mg guanidium-HCl (final concentration 6M) to each of the reaction tube. NaCl and imidazole was added to a final concentration of 300 mM and 10 mM respectively. Each sample was mixed with 100 µl Ni-NTA agarose beads and rotated O.N. at 4°C. Beads were collected by centrifugation and washed twice with 500 µl wash buffer 1 (50 mM Tris-HCl pH 7.6, 300 mM NaCl, 10 mM imidazole, 6mM Guanidinium-HCl, 0.1% NP-40, 5mM β-mercaptoethanol) and 4 times with 500 µl PNK buffer (50mM Tris-HCl pH 7.6, 10 mM MgCl_2_, 0.5% NP-40, 10 mM β-mercaptoethanol). 80 µl of PNK buffer containing 4U Thermosensitive Alkaline Phosphatase (TSAP) (Promega) and 2µl RNasin® (Promega) was added to the beads and incubated for 60 min at 37°C. Beads were washed once with 500 µl 1X Wash Buffer (same as above) and 3 times with 500 µl PNK buffer. Eighty µl of 1X PNK containing 2µl (20U) of T4 PNK (New England Biolabs), 2µl (80U) RNasin and 20 µCi γ32P-ATP (PerkinElmer) were added to the beads pellet and incubated 60 min at 37°C. Beads were washed once with 500 µl Wash 1 followed by 3 washes with 500 µl PNK buffer, and eluted with 100 µl elution buffer (50 mM Tris-HCl pH7.6, 50 mM NaCl, 200 mM imidazole, 0.1% NP-40, 5mM β-mercaptoethanol) for 5min at room temperature.

### Electrophoretic Mobility Shift Assay (EMSA) (gel-shift)

Purified Xrn1 was diluted with binding buffer (50 mM Tris-HCl, pH 7.4, 150 mM NaCl, 2 mM DTT, 1mM EDTA) in a total volume of in 10 μl. 5’-end labelled-probe^50^was heated at 95°C for 1 min, and snap chill in wet ice for 5 min. 100 CPS of radiolabelled probe, in a binding buffer, was added to each reaction tubes. The mixture was incubated for 30 min at room-temperature. One μl of loading buffer (80% glycerol + bromophenol blue + Xylene-Cyanol) was added and samples were fractionated by 6% non-denaturing polyacrylamide gel in 0.5 x TBE buffer. The gel was run for 3 hrs at 200 V in 0.5 X TBE in the cold-room. The gel was exposed to X-ray film at −70°C.

### Crosslinking and Analysis of cDNA (CRAC)

CRAC experiment and its analyses were performed as previously described^51–54^. Briefly, optimally growing cells carrying appropriate variations of HTP-tagged Xrn1 or Tail were UV crosslinked and harvested. RNA-protein complexes were captured tandemly on IgG sepharose and Ni-NTA followed by partial RNase digestion using RNase-IT. After ligation of sequencing adaptors, cDNA libraries were prepared by reverse transcription and PCR amplification followed by Illumina deep sequencing. Following the collection of *.fastq files, adapter trimming was done by flexbar^53, 54^. RA3 (5’-TGGAATTCTCGGGTGCCAAGG-3’) and RA5 (5’-GTTCAGAGTTCTACAGTCCGACGATCNNNNNAGC-3’) adapter of Illumina TruSeq are selected as input in the flexbar run. For further sequence processing and statistical analysis, pyCRAC packages were used (https://git.ecdf.ed.ac.uk/sgrannem/pycrac). pyFastqDuplicateRemover.py, a program which removes duplicate from the flexbar trimmed files were used for sequence processing (pyCRAC package). Next, Bowtie2 v2.4.2 was used to align trimmed sequence on R64-1-1 reference genome (https://uswest.ensembl.org/Saccharomyces_cerevisiae/Info/Annotation) and *.sam output format was produced for the downstream analyses. pyReadCounters.py program (from the same pyCRAC package) was used to calculate the RPKM values of genes.

### Genomic run-on

Genomic Run-On (GRO) was done in three biological replicates essentially as decribed^55, 56^. Exponentially growing yeast cells (5x10^8^ cells at OD_600_= 0.5) were taken for each run-on reaction. Another aliquot of the same cells was used directly for RNA extraction, which subsequently was used for cDNA synthesis using oligo-dT as the primer and ^33^P-dCTP to determine mRNA abundance (RA). GRO samples provided nascent transcription rates (nTR) for every yeast gene. Values were normalized to the average cell volume (the median of the population measured by a Coulter Counter device), although this correction had very little effect since cell volume of all the studied strains was similar. Whole RNA polymerase II TR was obtained by summing up all individual genes TR data. RAs were obtained from the hybridization of labelled cDNA onto nylon filters. Total mRNA concentration in yeast cells was determined by quantifying polyA + in total RNA samples by oligo-dT hybridization of a dot-blot following the protocol described ^57^and dividing by average cell volume. mRNA half-lives (HLs), in arbitrary units, were obtained by dividing individual RA values by TR ones.

### mRNA decay assay

mRNA decay assay Was performed as described previously ^10, 47, 58, 59^. To block transcription of non-stress genes, optimally proliferating cells (∼1 × 10^7^cells/ml) were shifted rapidly from 30°C to 42°C by shaking for 55 s at 70°C and then incubated at 42°C. Samples were taken at various time points post-temperature shift and mRNA levels were determined by Northern blot hybridization as described previously.

### Cell-cycle analysis

Cells were proliferated in SD medium at 30°C and allowed to enter stationary phase (SP). Following entry, cells were incubated in for 7 additional days. Cells were collected by centrifugation and were resuspended in fresh medium at 10 fold smaller cell density, followed by incubation at 30°C. 25 ml of Cells were withdrawn at each time-point and concentrated 3-fold. Cells were sonicated to break the clumps. 300 µl of cells were transferred to microcentrifuge tube, centrifuged and resuspended in 700 µl ethanol (final 70% ethanol), incubated at room temperature for 2h. Fixed cells were centrifuged, and the pellet was washed 4 times with 1ml of 0.2 M Tris HCl (pH 7.5). Pellet was resuspended in 0.5 ml Tris buffer. 10 µl 0.5 M EDTA and 50 µl RNaseA (10 mg/ml), and rotated overnight at 37°C. Cells were centrifuged and resuspended in pepsin (5 mg/ml in HCl) and incubated for 2 h at 37°C. Cells were washed with 0.2 M Tris and eventually were suspended in 0.3 ml of 0.2 M Tris. Cells were diluted 1:10 in 1 ml 0.2M Tris pH 7.5, 1µM SYTOX ^60–62^ was added just before injection into the flow cytometer (Beckton-Dickinson FACScalibur). 10,000 ungated events for each run were recorded using FL1 (488 nm laser). The data were analysed by FCS Express (DeNovo software).

### Data analysis and Statistics

Fluorescence images were analyzed by ImageJ^48, 49^. Immunoblotting and northern-blotting band intensities were quantitated by TotalLab software. SwissModel^63^was used to model Xrn1 and Tail, PyMol (The PyMOL Molecular Graphics System, Version 1.2r3pre, Schrödinger, LLC) was used to visualize and manipulate the model and for overlaying of structures. To determine sequence conservation, sequence logo was generated, using the online WebLogo program^64^(http://weblogo.berkeley.edu/logo.cgi). Protein disorder was predicted in PONDR^65, 66^ (http://www.pondr.com/). Data was plotted and statistical calculation were done in Excel or R environment using ‘ggplot2’, ‘eulerr’, ‘ggpubr’ and ‘plyr’ packages. For assessing the difference between two groups, *p*-value was calculated using Student’s unpaired T-test or Wilcoxon rank sum test (Also known as Mann-Whitney U test) as indicated in the Figure legends.

## Acknowledgements

We thank Gal Haimovich, Juana Diez, Shubham Dashmukh and Orna Amster-Choder for critically reading the manuscript. We thank Philipp Hackert for help with the CRAC experiments. This work was supported by the Israel Science Foundation (1472/15) to MC and by the Deutsche Forschungsgemeinschaft (SFB1190 to MTB. and KEB) and the Heidenreich von Siebold program of the University Medical Center Göttingen (to KEB). It was supported in part at the Technion by a fellowship of the Israel Councle for Higher Education (to SC).

